# Prior vaccination with the rVSV-ZEBOV vaccine does not interfere with but improves the efficacy of postexposure antibody treatment in nonhuman primates exposed to Ebola virus

**DOI:** 10.1101/2020.04.01.018275

**Authors:** Robert W. Cross, Zachary A. Bornholdt, Abhishek N. Prasad, Joan B. Geisbert, Viktoriya Borisevich, Krystle N. Agans, Daniel J. Deer, Kevin Melody, Karla A. Fenton, Heinz Feldmann, Armand Sprecher, Larry Zeitlin, Thomas W. Geisbert

## Abstract

A replication-competent, vesicular stomatitis virus vaccine expressing the Ebola virus (EBOV) glycoprotein (GP) (rVSV-ZEBOV) was successfully used during the 2013-16 EBOV epidemic^1^. Additionally, chimeric and human monoclonal antibodies (mAb) against the EBOV GP showed promise in animals and EBOV patients when administered therapeutically^2-6^. Given the large number of at-risk humans being prophylactically vaccinated with rVSV-ZEBOV, there is uncertainty regarding whether vaccination would preclude use of antibody treatments in the event of a known exposure of a recent vaccinee. To model a worst-case scenario, we performed a study using rhesus monkeys vaccinated or unvaccinated with the rVSV-ZEBOV vaccine. One day after vaccination, animals were challenged with a uniformly lethal dose of EBOV. Five vaccinated animals and five unvaccinated animals were then treated with the anti-EBOV GP mAb-based therapeutic MIL77 starting 3 days postexposure. Additionally, five vaccinated macaques received no therapeutic intervention. All five macaques that were vaccinated and subsequently treated with MIL77 showed no evidence of clinical illness and survived challenge. In contrast, all five animals that only received the rVSV-ZEBOV vaccine became ill and 2/5 survived; all five macaques that only received MIL77 only also became ill and 4/5 survived. Enhanced efficacy of vaccinated animals that were treated with MIL77 was associated with delayed EBOV viremia attributed to the vaccine. These results suggest that rVSV-ZEBOV augments immunotherapy.

---

Outbreaks of filovirus disease have become increasingly difficult to manage due to increased connectivity in endemic regions coupled with inadequate public health infrastructures and lack of approved medical countermeasures including diagnostics, therapeutics, and vaccines. Due to the sporadic nature of these outbreaks, the development and efficacy testing of preventative vaccines and postexposure treatments has previously been limited to animal models, including nonhuman primates, in which complete protection from lethal EBOV challenge has been demonstrated^7-9^. The unprecedented magnitude of the 2013-16 West African EBOV epidemic offered a unique opportunity to assess the efficacy of some of the most promising medical countermeasures available at that time^7,8^. Notably, the rVSV-ZEBOV vaccine was shown to provide 100% efficacy (95% CI 68·9–100·0, p=0·0045) when used in Guinea in a ring vaccination, open-label, cluster-randomized Phase III clinical trial^1^. The same vaccine has reportedly shown similar levels of success on a larger scale in the current outbreak of EBOV in the Democratic Republic of Congo (DRC), having been administered to over 200,000 people^10^. Built on the successes of years of development and validation in the field during two major ebolavirus outbreaks, the rVSV-ZEBOV vaccine (licensed as Ervebo™) was recently approved for human use by both the US FDA and the European Union^11^. Due to the widespread deployment of the rVSV-ZEBOV vaccine within a hot zone and to medical professionals, vaccinated individuals may encounter high risk exposures to EBOV prior to the development of protective immunity. The post-vaccination window of susceptibility has raised questions and concerns around the use of EBOV therapeutic options for such cases. The most significant concern regards the impact of potentially detrimental interference resulting from co-administration of a vaccine displaying EBOV GP as an immunogen with therapeutic mAbs that target EBOV GP, currently the most effective postexposure EBOV interventions available for human use^2-6^.

Previous studies have shown that the rVSV-ZEBOV vaccine causes a transient viremia in nonhuman primates (NHP) between days 2 and 4 after vaccination^12,13^. The half-life of rVSV-ZEBOV GP antigen in tissues of vaccinated primates is unknown; however, the presence of rVSV-ZEBOV GP in tissues was only detected in 2/6 pigs at day 3 post vaccination^14^. Thus, either circulating rVSV-ZEBOV or expression of EBOV GP from rVSV-ZEBOV-infected tissues could interfere with subsequent administration of any mAb-based therapeutic targeting EBOV GP.

In order to address the potential issue of vaccine/therapeutic or therapeutic/vaccine interference we employed a uniformly lethal rhesus macaque model of EBOV infection^9^. In brief, sixteen animals were divided into three experimental groups (n=5/group) and one control animal. Animals in one group were given the rVSV-ZEBOV vaccine on day -1 and then received the anti-EBOV GP mAb therapeutic MIL77 on days 3, 6, and 9 at 20 mg/kg/dose after EBOV exposure. The MIL77 immunotherapeutic was selected based on availability and previous results in EBOV-challenged NHPs^5^. We reduced the dose of MIL77 from a therapeutically proven dose of 50 mg/kg to 20 mg/kg^5^ to deliver a dose on the margin of protection and accentuate any potential interference from the rVSV-ZEBOV vaccine. Animals in the second experimental group only received the rVSV-ZEBOV vaccine on day -1 and animals in the third experimental group were only treated with MIL77 on days 3, 6, and 9 post-infection (dpi) (20 mg/kg/dose). The single EBOV challenge control animal was not vaccinated or given mAb therapy and succumbed to disease 9 days postexposure. Importantly, 12 historical control rhesus macaques challenged via the same route with the same EBOV seed stock and target dose all succumbed 6 to 9 days after challenge (**Figure 1a**)^9,15^.

**Figure 1:**
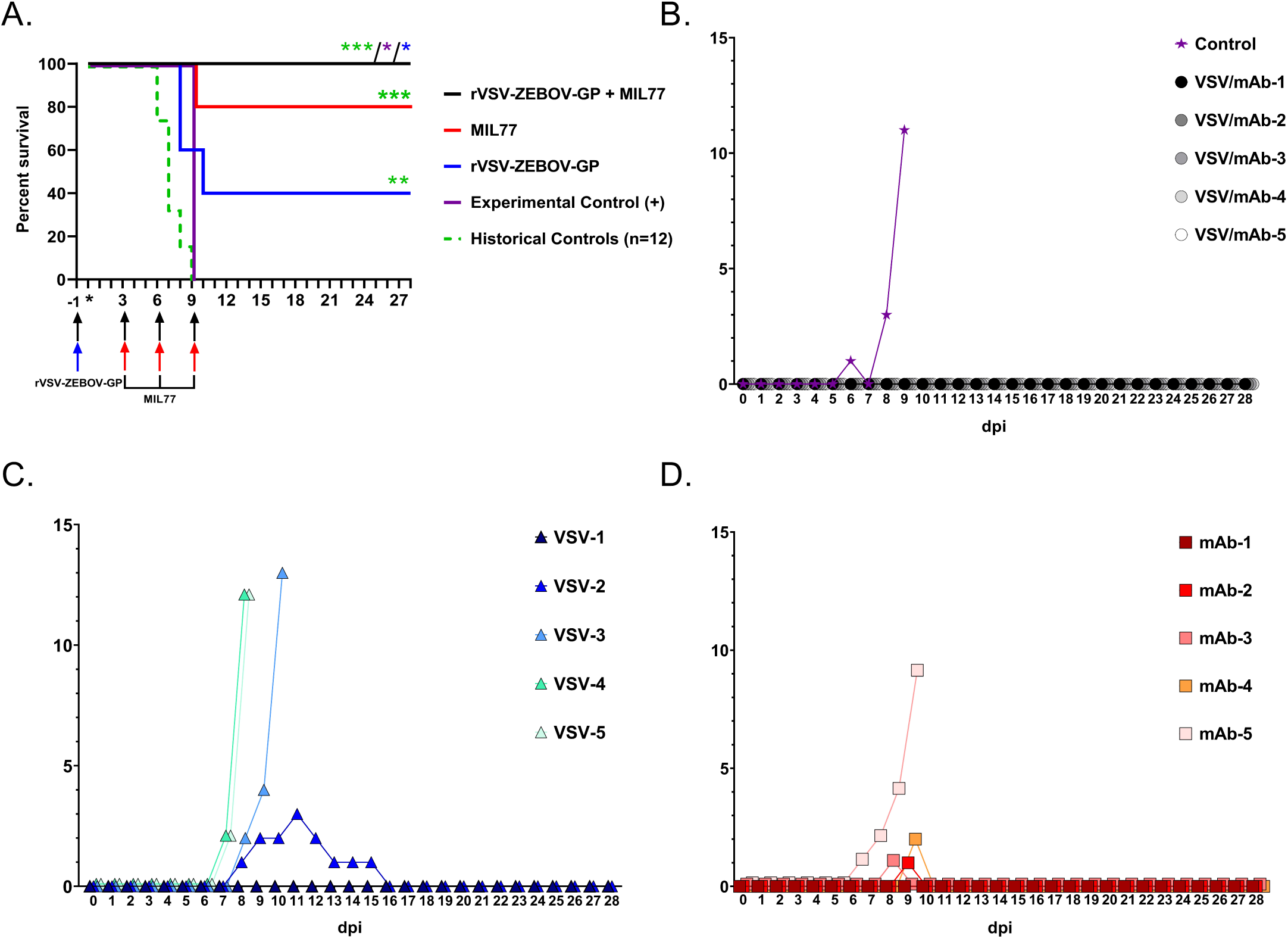
Survival and Clinical Score Outcomes: Animals were vaccinated/treated with rVSV-ZEBOV and/or MIL77 at the indicated day pre/post-infection. n=5 for all groups except experimental control (n=1). Historical control rhesus macaques infected with EBOV-Kikwit were included for statistical comparisons (n=12). Significance was measured using the Log-rank (Mantel-Cox) test. Colored asterisks denote statistical significance to the same colored group. p=0.0253 for rVSV-ZEBOV + MIL77 vs. experimental control cohort; p= 0.0486 for rVSV-ZEBOV + MIL77 vs. rVSV-ZEBOV cohort; p= 0.0003 for rVSV-ZEBOV + MIL77 vs. Historical Control cohort; p= 0.0072 for rVSV-ZEBOV vs. Historical Control cohort; p= 0.0009 for MIL77 vs. Historical Control cohort; all other comparisons were statistically insignificant. Arrows denote color-coded cohort. Asterisk indicates day of challenge (day 0). **(B-D)** Clinical illness scores for rVSV-ZEBOV + MIL77 (**B**), rVSV-ZEBOV (**C**), and MIL77 (**D**) treated rhesus macaques. For each panel, dashed lines indicate temperature (left y-axis) and solid lines indicate clinical score (right y-axis). Significance is graphically tiered as following: * = ≤ 0.05, ** = ≤ 0.005, and *** = ≤ 0.0005.

The unvaccinated/untreated control animal developed clinical symptoms of EBOV disease (EVD) beginning at 5 dpi (**Figure 1b, Supplementary Table 1**), and succumbed to disease at 9 dpi (**Figure 1a**). Animals that were either vaccinated day -1 with rVSV-ZEBOV or treated day 3, 6, and 9 dpi with MIL77 all developed clinical illness with 2/5 and 4/5 animals in each group surviving, respectively (**Figure 1c,d**). Notably, all five animals that were vaccinated day -1 with rVSV-ZEBOV and subsequently treated on days 3, 6, and 9 with MIL77 survived to the study endpoint (28 dpi) without developing any clinical signs of EVD. There was a significant difference in survival between the rVSV-ZEBOV + MIL77 treated group and the control animal (p = 0.0253, Mantel-Cox log-rank test), and between the rVSV-ZEBOV + MIL77 and rVSV-ZEBOV treatment groups (p=0.0486, Mantel-Cox log rank test) (**Figure 1a**). No significant difference was detected between the experimental control animal in this study and historical control (HC) rhesus macaques (N=12). However, differences were detected when comparing the experimental treatment groups with the HC NHPs (p= 0.0003 for rVSV-ZEBOV + MIL77 vs. HC; p= 0.0072 for rVSV-ZEBOV vs. HC; p= 0.0009 for MIL77 vs. HC).

There were notable differences in clinical pathology and the course of EVD between the experimental treatment groups. A single animal (VSV/mAb-1) in the dual-treatment group developed fever 1 dpi, which was most likely vaccine associated, whereas the control animal and 4/5 animals each in the vaccination only and mAb treatment only groups developed fever 6-9 dpi, which coincided with the appearance of other signs of EVD (**Figure 1b-d, Supplementary Tables 2,3**). Post-mortem pathological findings in the control animal and vaccinated or treated animals that succumbed was consistent with previous reports of EVD in macaques (**Figure 3d, h, l, p, t, and x)**^16,17^. Animals surviving challenge, including all animals in the rVSV-ZEBOV + MIL77 treatment group, exhibited no significant gross or histopathological findings (**Figure 3a-c, e-g, i-k, m-o, q-s, u-w)**.

**Figure 2:**
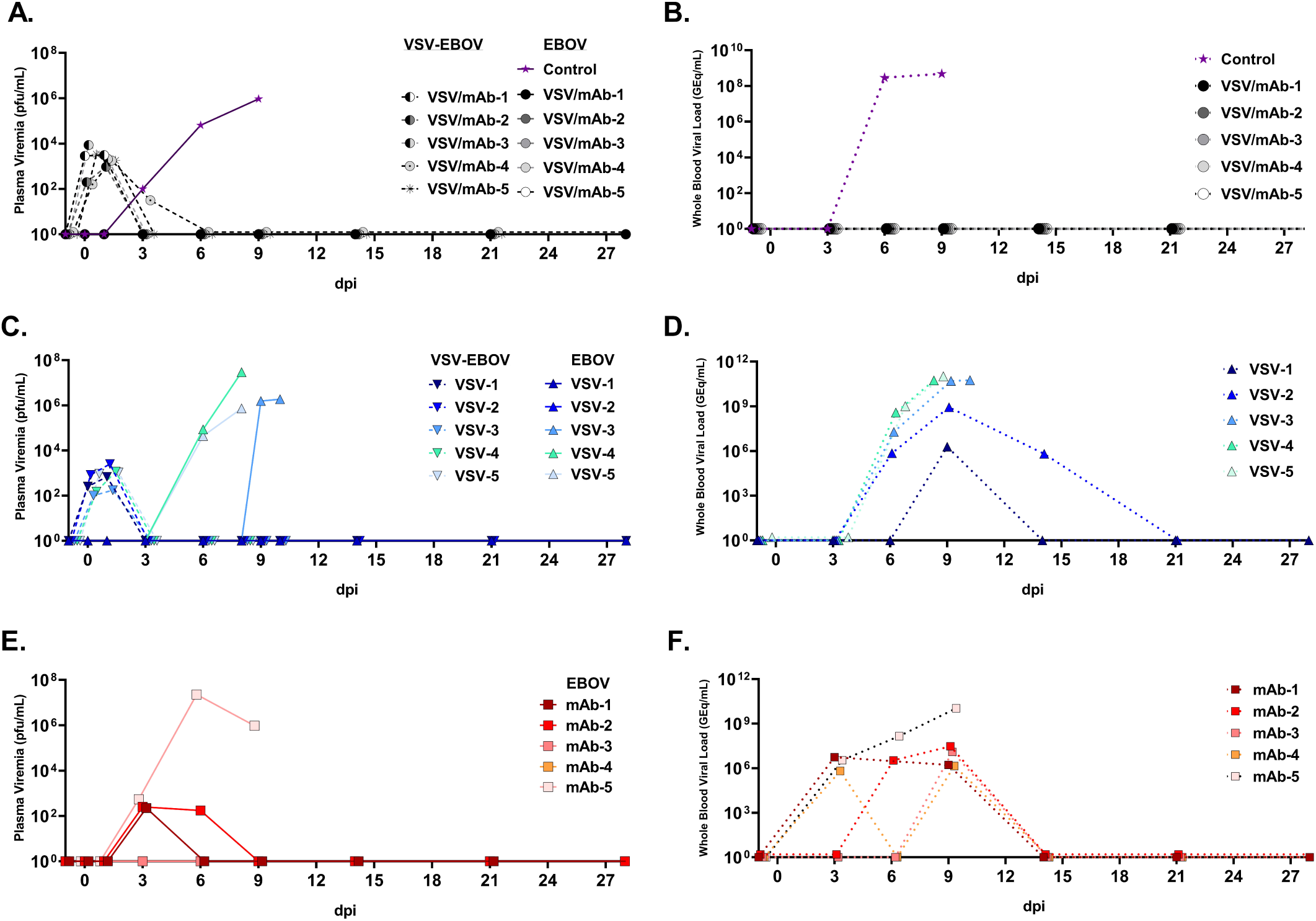
Circulating infectious virus and vRNA: Plasma viremia and vRNA content in whole blood of rVSV-ZEBOV + MIL77 (A. & B.), rVSV-ZEBOV (C. & D.), and MIL77 (E & F.). Limit of detection for infectious virus assays was 25 pfu/ml.

**Figure 3:**
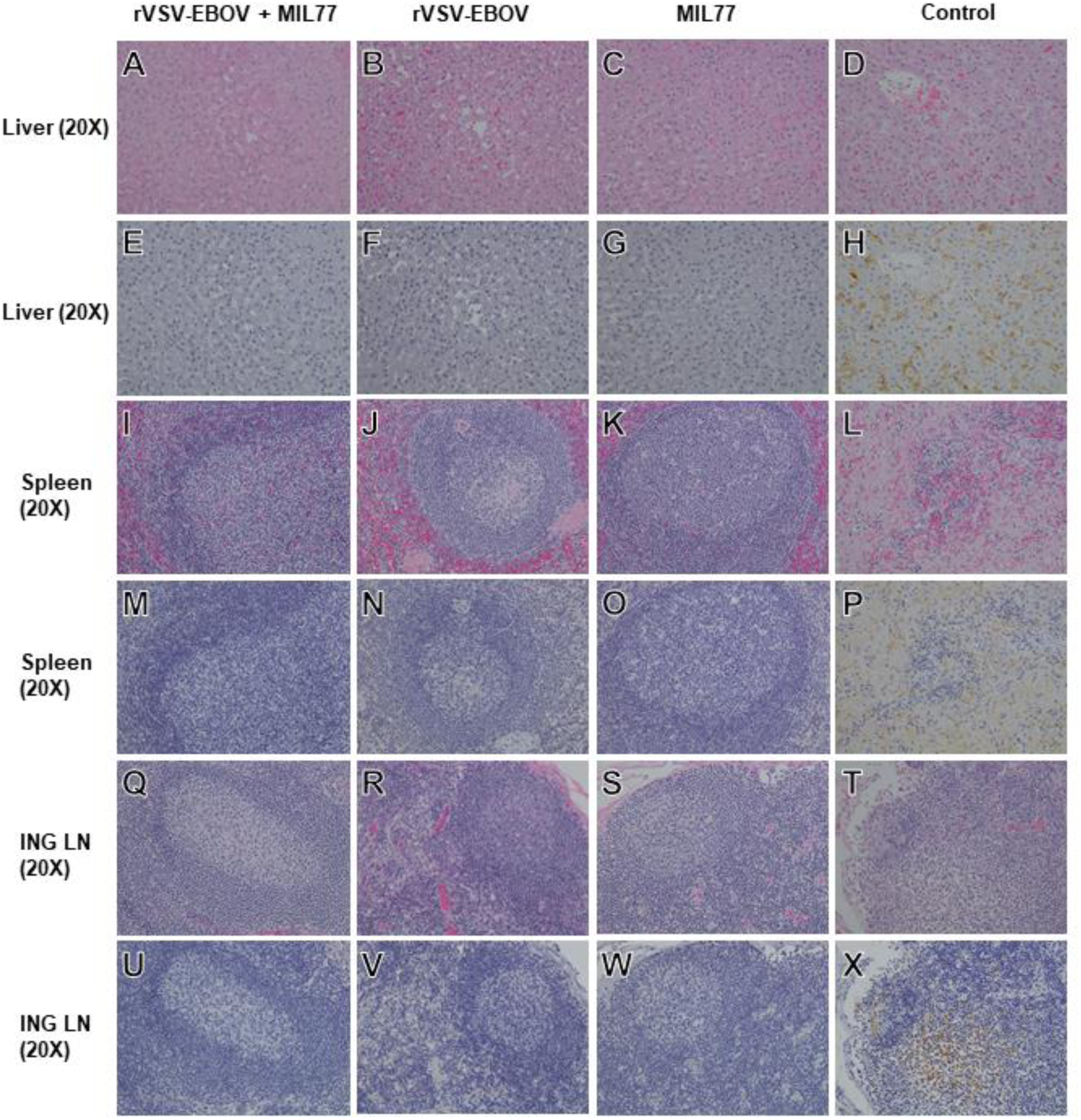
Hematoxylin/eosin and immunohistochemical staining of tissues from EBOV challenged rhesus macaques. Representative H&E-stained tissue specimens (A,B,C,D,I,J,K,L,Q,R,S,T) and IHC antibody labeled tissue specimens (E,F,G,H,M,N,O,P,U,V,W,X). ING LN=inguinal lymph node. For IHC images, EBOV antigen staining (VP40 protein), if present, is shown in brown. For the rVSV-EBOV + MIL77 cohort, images were collected from subject #VSV/mAb-2, and are representative of all animals in the cohort. For the rVSV-EBOV cohort, images were collected from subject #VSV-1, and are representative of all animals in the cohort surviving challenge. For the MIL77 cohort, images were collected from subject #mAb-4, and are representative of all animals in the cohort surviving challenge. Images from a historical control animal, subject #H-7, are representative of what was observed in the other historical controls and the control animal for this study. Animals from the rVSV-EBOV and MIL77 only cohorts which succumbed to disease exhibited histological lesions and antigen labeling comparable to control animals. All images were captured at 20X magnification.

Infectious rVSV-ZEBOV was detected by plaque assay up to 2 days post-vaccination in all vaccinated animals except one (VSV/mAb-4) which had low level (1.4 log_10_ pfu/ml) viremia on day 4 post-vaccination. Detection of circulating EBOV genomic RNA (vRNA) and infectious virus was performed by RT-qPCR and plaque assay titration, respectively. Consistent with historical controls challenged with the same EBOV seed stock, the experimental control animal had 2 log_10_ pfu/ml of infectious EBOV by 3 dpi and 8.46 log_10_ GEq of vRNA by 6 dpi, which then peaked on 9 dpi at euthanasia for both detection methods (**Figure 2a, Supplementary Figure 1)**. The vaccine only group had detectable EBOV vRNA by 6 dpi in 4/5 animals and by 9 dpi in the remaining animal (**Figure 2d**). Infectious EBOV was detected in the same group by 6 dpi in 2/5 animals with peak viral titers comparable to both the experimental and HC animals (**Figure 2c, Supplementary Figure 1**). In the MIL77 only group, EBOV vRNA was detected by 3 dpi in 3/5 animals and in 5/5 by 9 dpi. Likewise, infectious EBOV was detected in 3/5 animals by 3 dpi (**Figure 2e**). Infectious virus became undetectable in two surviving animals by 9 dpi, while the third animal was euthanized at 9 dpi. In stark contrast, none of the rVSV-EBOV + MIL77 animals had detectable circulating infectious EBOV or EBOV vRNA at any point postexposure (**Figure 2a, b**), consistent with the total lack of clinical scoring in this group.

We used ELISA-based detection to estimate total host derived anti-VP40 IgM and IgG as well as anti-GP IgM (**Figure 4**). Notably, the rVSV-ZEBOV + MIL77 group (3/5) and the rVSV-ZEBOV animals that survived (2/5) had detectable IgM to VP40 and GP by 6 dpi (**Figure 4a,b,d,e**), yet the IgM responses in the MIL77 group were at or below the limit of detection for the assays (**Figure 4c,f**). All animals from the rVSV-ZEBOV + MIL77 and any surviving animals from the rVSV-ZEBOV or MIL77 groups had clear evidence of circulating IgG antibodies against VP40 at 9 dpi through the end of the study (**Figure 4g-i**). Circulating amounts of MIL77 were equivalent across both mAb treated groups through the acute phase of disease; however, a slightly faster rate of decay was noted in 2/5 of the MIL77 only animals at study endpoint (**Figure 4j**).

**Figure 4:**
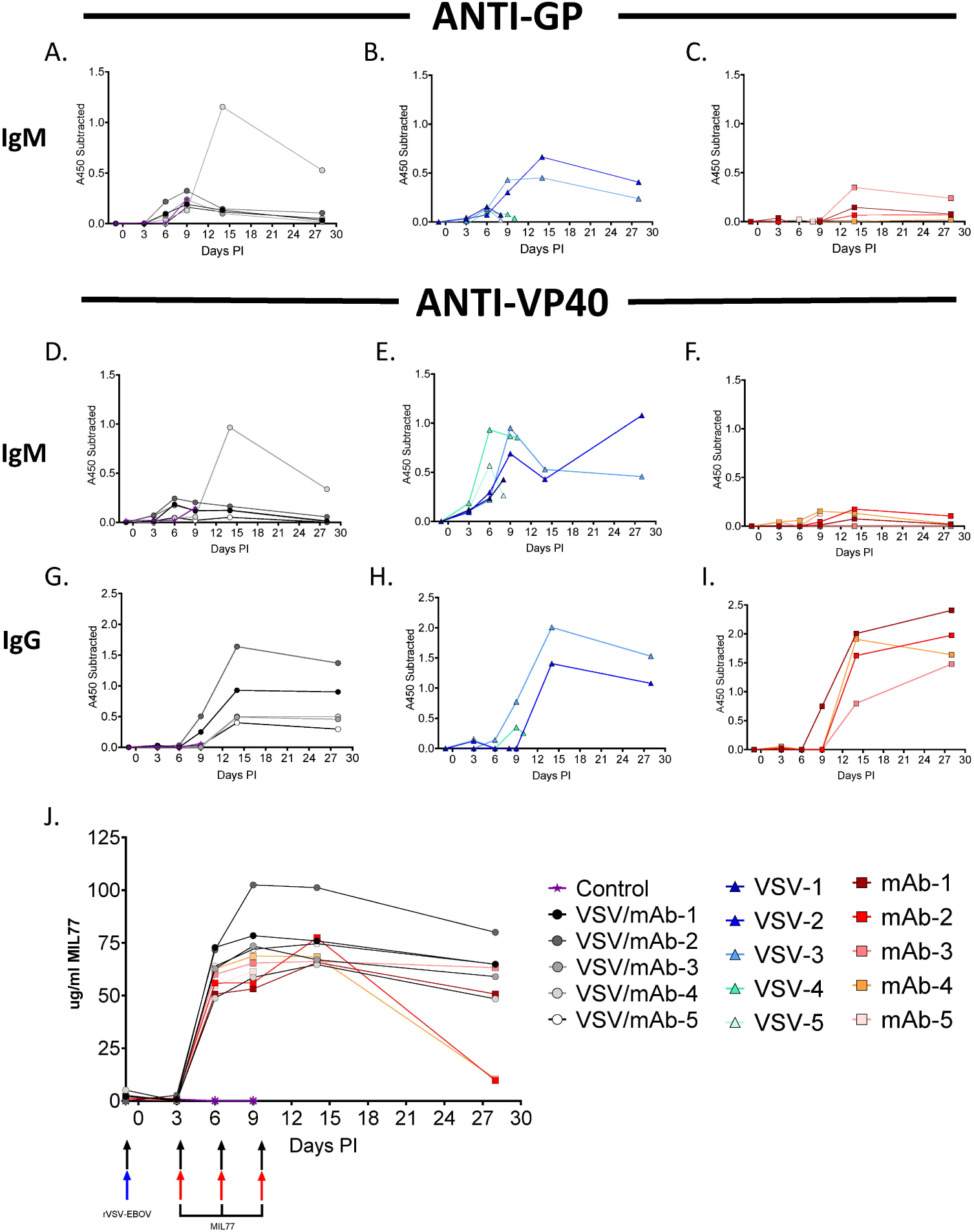
Circulating host and therapeutic antibodies in EBOV challenged and treated rhesus macaques. Circulating anti-GP IgM from rVSV-EBOV + MIL77 (**A**.), rVSV-EBOV (**B**.), and MIL77 (**C**.) groups. Circulating anti-VP40 IgM from rVSV-EBOV + MIL77 (**D**.), rVSV-EBOV (**E**.), and MIL77 (**F**.) groups. Circulating anti-VP40 IgG from rVSV-EBOV + MIL77 (**G**.), rVSV-EBOV (**H**.), and MIL77 (**I**.) groups. ELISA data are represented as mean technical replicates (n=2) where change in absorbances at 450nM from subtracted from baseline at day -1 (A-I.). Circulating MIL77 concentrations for the rVSV-EBOV + MIL77 and MIL77 groups are represented over time (**J**.). Significance of concentration differences between rVSV-EBOV + MIL77 and MIL77 only groups (P&≤ 0.001 = ***) were determined with two tailed t-tests using Holm-Sidak correction for multiple comparisons.

The 2013-16 West African and current EBOV epidemic in the DRC, both of previously unprecedented proportion, have demonstrated the critical need for efficacious medical countermeasures. However, the potential for deleterious interference between different modes of treatment presents a possible barrier to the development and approval of protocols utilizing a combinatorial approach. Studies in NHPs investigating protection by the rVSV-ZEBOV vaccine when administered as a postexposure intervention have demonstrated only partial protection suggesting that additional postexposure countermeasures may be necessary^18,19^. Indeed, seroconversion offering protective immunity does not occur before 3 days post-vaccination in NHPs^20^, and the same is likely true in humans^21^. Given that all current candidate postexposure mAb therapeutics in clinical trials target the EBOV GP, which is also the antigenic immunogen displayed by the rVSV-ZEBOV vaccine vector, significant concern exists regarding the potential for interference between these types of products. Accordingly, we performed a narrowly focused study utilizing rhesus monkeys to model a scenario likely occurring during the current outbreak in DRC; namely, high-risk exposure to EBOV in individuals recently vaccinated with rVSV-ZEBOV. To assess the potential contraindication of subsequent mAb treatment, we treated a cohort of vaccinated animals with the MIL77 mAb cocktail at days 3, 6, and 9 dpi. Surprisingly, instead of interference, we observed clear therapeutic benefit, where animals vaccinated before EBOV challenge and then subsequently treated postexposure were afforded complete protection without any observable clinical disease. In contrast, animals that received vaccination only or mAb treatment only displayed significant signs of clinical EVD, and in the case of the vaccine only group, limited protection.

It has recently been demonstrated that induction of potent innate immune effector mechanisms occurs in the context of rVSV-ZEBOV vaccination^22^. Indeed, others have shown modest widening of the therapeutic window upon administration of exogenous interferon-alpha modalities, although neither approach was enough to induce protection ^23,24^. While the precise mechanism is unclear, the scenario presented here suggests reduction of circulating infectious EBOV complements the induction of vaccine-induced EBOV immunity, ultimately reducing morbidity and likely contributing to survival. Of note, a precedent for a tandem approach of vaccination and mAb treatment for postexposure treatment is the standard protocol for rabies virus exposure, which recommends both vaccination with the inactivated rabies virus vaccine and treatment with human rabies immunoglobulin^25^. Our study suggests that a similar approach to treatment may be appropriate for high-risk EBOV exposure and that mAb therapy post-vaccination may improve clinical outcome in recently vaccinated individuals.

## Methods

### Challenge virus. Zaire ebolavirus

(EBOV) isolate 199510621 (strain Kikwit) originated from a 65-year-old female patient who had died on 5 May 1995. The study challenge material was from the second Vero E6 passage of EBOV isolate 199510621. Briefly, the first passage at UTMB consisted of inoculating CDC 807223 (passage 1 of EBOV isolate 199510621) at a MOI of 0.001 onto Vero E6 cells. The cell culture fluids were subsequently harvested at day 10 post infection and stored at -80°C as ∼1 ml aliquots. Deep sequencing indicated the EBOV was greater than 98% 7U (consecutive stretch of 7 uridines). No detectable mycoplasma or endotoxin levels were measured (< 0.5 endotoxin units (EU)/ml).

### Nonhuman primate vaccination, challenge, and treatment

Sixteen healthy, filovirus-naive, adult (∼ 3.7 to 5.4 kg) Chinese origin rhesus macaques (*Macaca mulatta*; PreLabs) were randomized into three groups of five experimental animals each and a control group of one animal. Animals in two of the experimental animals were vaccinated by intramuscular (i.m.) injection of ∼ 2×10^7 PFU of the rVSV-ZEBOV GP vaccine based on the Mayinga strain^26^; this is the same dose used for humans^1^. Animals in the remaining experimental group as well as the control animal were not vaccinated. All 16 of the macaques were challenged one day after vaccination of the experimental groups by i.m. injection with a target dose of 1,000 PFU of EBOV strain Kikwit. Experimental animals in one of the vaccinated groups (rVSV-ZEBOV + MIL77) and one of the unvaccinated groups (MIL77 only) were treated by intravenous (i.v.) administration of ∼ 20 mg/kg of MIL77 on days 3, 6, and 9 after EBOV challenge while animals in the experimental unvaccinated group (rVSV-ZEBOV only) and the control animal were not treated. All 16 animals were given physical examinations, and blood was collected before vaccination (day -1), before virus challenge (day 0), and on days 1, 3, 6, 9, 14, 21, and 28 (study endpoint) after virus challenge. The macaques were monitored daily and scored for disease progression with an internal EBOV humane endpoint scoring sheet approved by the UTMB IACUC. UTMB facilities used in this work are accredited by the Association for Assessment and Accreditation of Laboratory Animal Care International and adhere to principles specified in the eighth edition of the Guide for the Care and Use of Laboratory Animals, National Research Council. The scoring changes measured from baseline included posture and activity level, attitude and behavior, food intake, respiration, and disease manifestations, such as visible rash, hemorrhage, ecchymosis, or flushed skin. A score of ≥ 9 indicated that an animal met the criteria for euthanasia.

### Detection of viremia

RNA was isolated from whole blood utilizing the Viral RNA mini-kit (Qiagen) using 100 µl of blood added to 600 µl of the viral lysis buffer. Primers and probe targeting the VP30 gene of EBOV were used for real-time quantitative PCR (RT-qPCR) with the following probes: EBOV, 6-carboxyfluorescein (FAM)-5= CCG TCA ATC AAG GAG CGC CTC 3=-6 carboxytetramethylrhodamine (TAMRA) (Life Technologies).Viral RNA was detected using the CFX96 detection system (Bio-Rad Laboratories, Hercules, CA) in one-step probe RT-qPCR kits (Qiagen) with the following cycle conditions: 50°C for 10 min, 95°C for 10 s, and 40 cycles of 95°C for 10 s and 57°C for 30 s. Threshold cycle (CT) values representing viral genomes were analyzed with CFX Manager software, and the data are shown as genome equivalents (GEq) per milliliter. To create the GEq standard, RNA from viral stocks was extracted, and the number of strain-specific genomes was calculated using Avogadro’s number and the molecular weight of each viral genome.

Virus titration was performed for rVSV-ZEBOV and for EBOV by plaque assay with Vero E6 cells from all plasma samples as previously described^26^. Briefly, increasing 10-fold dilutions of the samples were adsorbed to Vero E6 monolayers in duplicate wells (200 µL); the limit of detection was 25 PFU/mL.

### Hematology and serum biochemistry

Total white blood cell counts, white blood cell differentials, red blood cell counts, platelet counts, hematocrit values, total hemoglobin concentrations, mean cell volumes, mean corpuscular volumes, and mean corpuscular hemoglobin concentrations were analyzed from blood collected in tubes containing EDTA using a laser-based hematologic analyzer (Beckman Coulter). Serum samples were tested for concentrations of alanine aminotransferase (ALT), albumin, alkaline phosphatase (ALP), amylase, aspartate aminotransferase (AST), C-reactive protein (CRP), calcium, creatinine, gammaglutamyltransferase (GGT), glucose, total protein, blood urea nitrogen (BUN), and uric acid, and by using a Piccolo point-of-care analyzer and Biochemistry Panel Plus analyzer discs (Abaxis).

### Histopathology and immunohistochemistry

A partial necropsy was performed on all subjects. Tissue samples of major organs were collected for histopathologic and immunohistochemical examination, immersion-fixed in 10% neutral buffered formalin, and processed for histopathology as previously described^27,28^. For immunohistochemistry, specific anti-ZEBOV immunoreactivity was detected using an anti-ZEBOV VP40 primary antibody (IBT) at a 1:4000 dilution for 60 minutes. The tissues were processed for immunohistochemistry using the Thermo Autostainer 360 (ThermoFisher, Kalamazoo, MI). Secondary used was biotinylated goat anti-rabbit IgG (Vector Labs, Burlingame, CA #BA-1000) at 1:200 for 30 minutes followed by Vector Horseradish Peroxidase Streptavidin, R.T.U (Vector) for 30 min. Slides were developed with Dako DAB chromagen(Dako, Carpenteria, CA #K3468) for 5 minutes and counterstained with Harris Hematoxylin for 30 seconds. Non-immune rabbit IgG was used as a negative control.

### Detection of total IgM and IgG responses to Ebola VP40 and GP

ELISA plates were coated overnight at 4°C with 0.1 µg/mL of EBOV GP-TM (Integrated Biotherapeutics) or recombinant VP40 antigen coated plates (Zalgen) were used both of which were blocked for 2 hours prior to use. Serum samples were assayed using a 1:100 dilution in ELISA diluent (1% heat-inactivated fetal bovine serum, 1× phosphate-buffered saline, and 0.2% Tween-20). Samples were incubated for 1 hour at ambient temperature and then removed, and plates were washed. Wells then were incubated for 1 hour with goat anti-NHP IgM or IgG conjugated to horseradish peroxidase (Fitzgerald Industries International) at a 1:5000 dilution. Wells were washed and then incubated with tetramethylbenzidine substrate (KPL) (100 uL/well) and incubated for 10 minutes followed by stop solution (100 uL/well). Microplates are read at 450 nm with 650 nm subtraction with an OD450 nm cut-off of 0.069.

### Detection of Circulating MIL77 antibody

ELISA plates were coated overnight at 4°C with 0.1 µg/mL of mouse anti-human IgG (human CH2 domain with no cross-reactivity to rhesus macaque IgG; clone R10Z8E9; BioRad) and then blocked for 2 hours. The serum samples were assayed at 4-fold dilutions starting at a 1:100 dilution in ELISA diluent (1% heat-inactivated fetal bovine serum, 1× phosphate-buffered saline, and 0.2% Tween-20). Samples were incubated for 1 hour at ambient temperature and then removed, and plates were washed. Wells then were incubated for 1 hour with goat anti-human IgG conjugated to horseradish peroxidase (Fitzgerald Industries International) at a 1:20,000 dilution. Wells were washed and then incubated with tetramethylbenzidine substrate (KPL) (100 uL/well) and incubated for 10 minutes followed by stop solution (100 uL/well). Microplates are read at 450 nm with 650 nm subtraction with an OD450 nm cut-off of 0.071(Biotek Cytation system). mAb were quantified using Prism software, version 7.04 (GraphPad), to analyze sigmoidal dose-response (variable slope), using MIL77 as standard.

### Statistical analysis

Specific statistical tests are noted in the text and/or figure legends. All statistical analysis was performed in Graphpad 8.2.1.

### Study approval

The animal studies were performed at the Galveston National Laboratory, University of Texas Medical Branch at Galveston and were approved by the University of Texas Medical Branch Institutional Animal Care and Use Committee.

### Availability of data and materials

The datasets used and/or analyzed during the current study are available from the corresponding author on reasonable request.

## Acknowledgments

The authors would like to thank the UTMB Animal Resource Center for husbandry support of laboratory animals, Dr. Chad Mire for assistance with the animal studies, and Natalie Dobias for expert histology and immunohistochemistry support. We also thank Drs. Luis Branco and Matt Boisen of Zalgen for generously providing EBOV VP40 antigen coated plates for the ELISA work. We also thank Dr. Tina Parker for helpful discussions. This study was supported by the Department of Health and Human Services, National Institutes of Health grants U19AI109711 and U19AI142785 to T.W.G. and UC7AI094660 for BSL-4 operations support of the Galveston National Laboratory. Partial funding was provided through the Intramural research program, NIAID/NIH to H.F. Opinions, interpretations, conclusions, and recommendations are those of the authors and are not necessarily endorsed by the University of Texas Medical Branch or the National Institutes of Health.

## Author contributions

Z.A.B., A.S., L.Z., H.F., and T.W.G. conceived and designed the study. J.B.G., R.W.C., D.J.D., and T.W.G. performed the NHP vaccination, infection, and treatment experiments and conducted clinical observations of the animals. V.B. and K.N.A. performed the clinical pathology assays. J.B.G. and V.B. performed the EBOV infectivity assays. K.N.A. performed the PCR assays. R.W.C. and K.M. performed the ELISA assays. A.N.P., Z.A.B., J.B.G., R.W.C., V.B., K.N.A., D.J.D., K.M., K.A.F., A.S., L.Z., H.F., and T.W.G. analyzed the data. K.A.F. performed histological and immunohistochemical analysis of the data. A.N.P., R.W.C, K.A.F., and T.W.G. wrote the paper. Z.A.B., A.S., L.Z., and H.F. edited the paper. All authors had access to all of the data and approved the final version of the manuscript.

## Competing interests

Z.A.B. and L.Z. are employees of Mapp Biopharmaceutical. H.F. and T.W.G. are listed on patents related to Ebola vaccines.

## Figure Legends

**Supplementary Figure 1: Historical control rhesus macaques infected with EBOV-Kikwit:** A. Infectious virus in circulation. Limit of detection is 25 pfu/ml. B. vRNA content in whole blood.

**Supplementary Table 1. Clinical description and outcome of rVSV-EBOV-GP vaccinated/MIL77 treated and control rhesus macaques following EBOV challenge:** Days after EBOV challenge are in parentheses. Lymphopenia, granulopenia, monocytopenia, and thrombocytopenia are defined by a ≥ 35% drop in numbers of lymphocytes, granulocytes, monocytes, and platelets, respectively. Leukocytosis, monocytosis, and granulocytosis are defined by a two-fold or greater increase in numbers of white blood cells over base line. Fever is defined as a temperature more than 2.5 °F over baseline, or at least 1.5 °F over baseline and ≥ 103.5 °F. Hypothermia is defined as a temperature ≤ 3.5°F below baseline. Hyperglycemia is defined as a two-fold or greater increase in levels of glucose. Hypoglycemia is defined by a ≥ 25% decrease in levels of glucose. Hypoalbuminemia is defined by a ≥ 25% decrease in levels of albumin. Hypoproteinemia is defined by a ≥ 25% decrease in levels of total protein. Hypoamylasemia is defined by a ≥ 25% decrease in levels of serum amylase. Hypocalcemia is defined by a ≥ 25% decrease in levels of serum calcium. (ALT) alanine aminotransferase, (AST) aspartate aminotransferase, (ALP) alkaline phosphatase, (CRE) Creatinine, (CRP) C-reactive protein, (Hct) hematocrit, (Hgb) hemoglobin

**Supplementary Table 2. Clinical description and outcome of rVSV-EBOV-GP vaccinated rhesus macaques following EBOV challenge:** Days after EBOV challenge are in parentheses. Lymphopenia, granulopenia, monocytopenia, and thrombocytopenia are defined by a ≥35% drop in numbers of lymphocytes, granulocytes, monocytes, and platelets, respectively. Leukocytosis, monocytosis, and granulocytosis are defined by a two-fold or greater increase in numbers of white blood cells over base line. Fever is defined as a temperature more than 2.5 °F over baseline, or at least 1.5 °F over baseline and ≥ 103.5 °F. Hypothermia is defined as a temperature ≤3.5°F below baseline. Hyperglycemia is defined as a two-fold or greater increase in levels of glucose. Hypoglycemia is defined by a ≥ 25% decrease in levels of glucose. Hypoalbuminemia is defined by a ≥ 25% decrease in levels of albumin. Hypoproteinemia is defined by a ≥ 25% decrease in levels of total protein. Hypoamylasemia is defined by a ≥ 25% decrease in levels of serum amylase. Hypocalcemia is defined by a ≥ 25% decrease in levels of serum calcium. (ALT) alanine aminotransferase, (AST) aspartate aminotransferase, (ALP) alkaline phosphatase, (CRE) Creatinine, (CRP) C-reactive protein, (Hct) hematocrit, (Hgb) hemoglobin

**Supplementary Table 3. Clinical description and outcome of MIL77 treated rhesus macaques following EBOV challenge:** Days after EBOV challenge are in parentheses. Lymphopenia, granulopenia, monocytopenia, and thrombocytopenia are defined by a ≥35% drop in numbers of lymphocytes, granulocytes, monocytes, and platelets, respectively. Leukocytosis, monocytosis, and granulocytosis are defined by a two-fold or greater increase in numbers of white blood cells over base line. Fever is defined as a temperature more than 2.5 °F over baseline, or at least 1.5 °F over baseline and ≥ 103.5 °F. Hypothermia is defined as a temperature ≤ 3.5°F below baseline. Hyperglycemia is defined as a two-fold or greater increase in levels of glucose. Hypoglycemia is defined by a ≥ 25% decrease in levels of glucose. Hypoalbuminemia is defined by a ≥ 25% decrease in levels of albumin. Hypoproteinemia is defined by a ≥ 25% decrease in levels of total protein. Hypoamylasemia is defined by a ≥ 25% decrease in levels of serum amylase. Hypocalcemia is defined by a ≥ 25% decrease in levels of serum calcium. (ALT) alanine aminotransferase, (AST) aspartate aminotransferase, (ALP) alkaline phosphatase, (CRE) Creatinine, (CRP) C-reactive protein, (Hct) hematocrit, (Hgb) hemoglobin.

